## Supplemental Material for "A green lifetime biosensor for calcium that remains bright over its full dynamic range"

**Contents:**

**1 supplemental Note**

**15 supplemental Figures**

**2 supplemental Tables**

**Supplemental Note 1**

To determine the fraction of G-Ca-FLITS in the calcium-bound state for the lifetime calibration, the intensity contribution each state needs to be determined for the specific microscope settings used for the calibration. To this end, we applied a theoretical correction of both the extinction coefficient and the quantum yield to obtain a relative brightness per state (Figure S10).

The extinction coefficient for the calcium-bound and calcium-free state was calculated over the full absorption spectrum by considering:

a. The normalized brightness of the excitation light (LED) $l\left( \lambda\right)$
b. The normalized transmission of the excitation filter $ExF\left( \lambda\right)$
c. The normalized reflection of the dichroic mirror $DichrR\left( \lambda\right)$
d. The normalized excitation spectra, for the corresponding state of the sensor $F_{ex}\left( \lambda\right)$
e. The maximal extinction coefficient $\varepsilon_{max}$

The effective extinction coefficient ($\varepsilon_{eff}$) is calculated as:

$$\varepsilon_{eff}=\varepsilon_{max}\frac{\int l\left( \lambda\right)\cdot ExF\left( \lambda\right)\cdot DichrRF\left( \lambda\right)\cdot F_{ex}\left( \lambda\right)d\lambda}{\int l\left( \lambda\right)\cdot ExF\left( \lambda\right)\cdot DichrR\left( \lambda\right)d\lambda}$$

The relative excitation efficiency ($e_{rel}$) with the deployed excitation source and filters is defined as:

$$e_{rel}=\frac{\varepsilon_{eff}}{\varepsilon_{max}}=\frac{\int l\left( \lambda\right)\cdot ExF\left( \lambda\right)\cdot DichrR\left( \lambda\right)\cdot F_{ex}\left( \lambda\right)d\lambda}{\int l\left( \lambda\right)\cdot ExF\left( \lambda\right)\cdot DichrR\left( \lambda\right)d\lambda}$$

The corrected emission spectra ($F_{em}\left( \lambda\right)$) of both the calcium-free and the calcium-bound state were corrected by considering:

a. The normalized transmission of the dichroic mirror $DichrT\left( \lambda\right)$
b. The normalized transmission of the emission filter $EmF\left( \lambda\right)$
c. The normalized sensitivity of the camera $S\left( \lambda\right)$
d. The quantum yield of the sensor QY

From these the relative sensitivity ($S_{rel}$) for detection of the sensor can be computed:

$$S_{rel}=\frac{\int EmF\left( \lambda\right)\cdot DichrTF\left( \lambda\right)\cdot S\left( \lambda\right)\cdot F_{em}\left( \lambda\right)d\lambda}{\int F_{em}\left( \lambda\right)d\lambda}$$

Consequently, the relative overall measured brightness ($B_{rel}$) for a specific combination of optical components is defined as:

$$B_{rel}=e_{rel}\varepsilon_{max}S_{rel}\mathrm{QY}$$

Applying this calculation to the spectra of G-Ca-FLITS using the settings throughout this manuscript (LED centered around 446 nm, excitation filter ff01-448/20, dichroic di02-r488-25, emission filter ff01-520/35), result in an $\varepsilon_{eff}$ of 15900 and 21300 M^-1^cm^-1^ for the calcium-free and -bound states, respectively. The $S_{rel}$ is 0.41 for both states. Combining $\varepsilon_{eff}$ and $S_{rel}$with the determined QY and $\varepsilon_{max}$, results in a $B_{rel}$ of 2710 and 2270 M^-1^cm^-1^ for the calcium-free and -bound states, respectively. Therefore, the intensity contribution of the calcium-free state is 1.2-fold higher than the calcium-bound state.

Using the same calculation for the sensor Tq-Ca-FLITS with suitable settings (LED centered around 446 nm, excitation filter ff01-448/20, dichroic di02-r442-25, emission filter ff01-482/25), results in a 3.6-fold higher intensity contribution of the calcium-bound state compared to the calcium-free state. This is almost identical to the ratio of 3.51 measured and reported previously, where the same setup and settings were applied (van der Linden et al., 2021, doi:10.1038/s41467-021-27249-w).

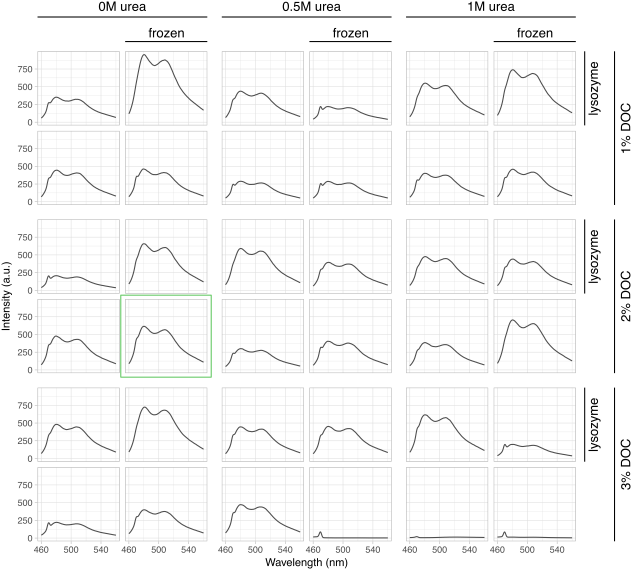

**Figure S1. Influence of different components of a bacterial lysis protocol.** Emission spectra of bacterial lysates expressing Tq-Ca-FLITS are shown. Various concentrations of urea, DOC and lysozyme were used in the lysis buffer. Each buffer also contained 50mM Tris-HCl (pH 8.0). The influence of a freeze/thawing step is also tested, indicated by ‘frozen’. Each condition was measured once. The condition that was selected for the screening is indicated with a green box.

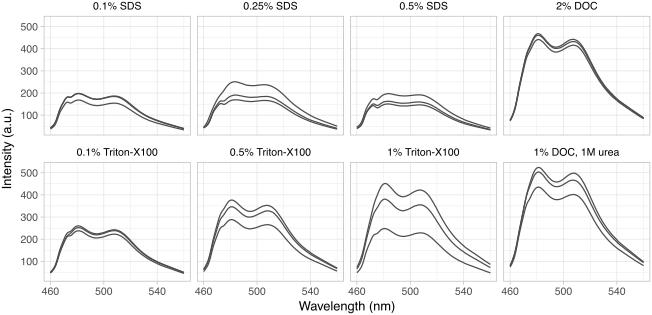
**Figure S2. Influence of different detergents on a bacterial lysis protocol.** Emission spectra of bacterial lysates expressing Tq-Ca-FLITS are shown. Various concentrations of SDS and Triton-X100 were tested and compared to a buffer with 2% DOC and a buffer with 1% DOC and 1M urea (the original bacterial lysis buffer^21^. Each buffer also contained 50mM Tris-HCl (pH 8.0). Each condition was measured in three times.

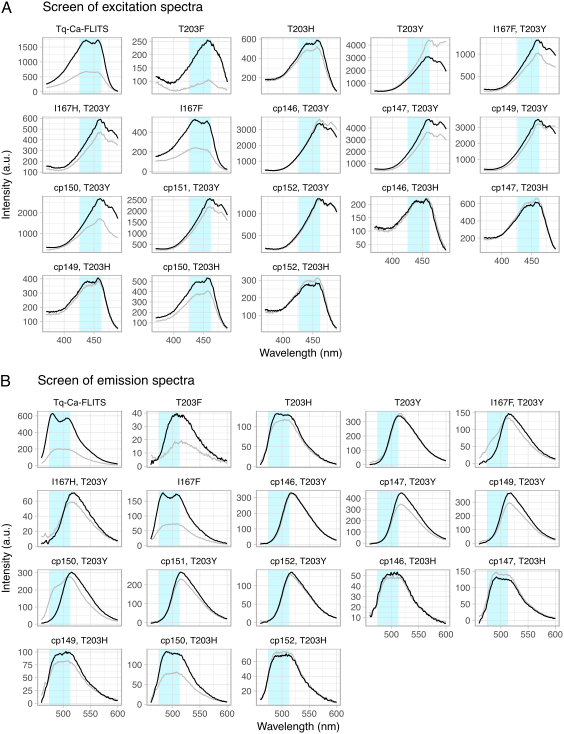

**Figure S3. Excitation (A) and emission spectra (B) of screened variants in the search for a red-shifted calcium sensor based on Tq-Ca-FLITS.** The spectra were measured in bacterial lysates, with addition of 0.1 mM CaCl_2_ (black line) or 9.5 mM EDTA (gray line). Sensor variants are indicated by their circular permutation and mutations in the FP in the sensor. For example, ‘cp146’ has from N- to C-terminus the following domains: calmodulin binding peptide M13, amino acids 146-283 of mTq2, a flexible GGSGG linker, amino acids 1-145 of mTq2, CaM (calmodulin). If no circular permutation position is indicated, the mutation displayed is done on Tq-Ca-FLITS. ‘T203Y’ indicates a T203Y mutation on the mTq2 in the sensor. The light blue shading is used as a reference for excitation and emission peaks of Turquoise.

```
             1                     21                            50
Tq-Ca-FLITS  MVDSSRRKWN KAGHAVRAIG RLSSPVVVYI TADKQKNGIK ANFKIRHNIE
G-Ca-FLITS   MVDSSRRKWN KAGHAVRAIG RLSSPVAVYI TADKQKNGIK ANFKIRHNIE

             51                    71                           100
Tq-Ca-FLITS  DGGVQLADHY QQNTPIGDGP VLLPDNHYLS TQSKLSKDPN EKRDHMVLLE
G-Ca-FLITS   DGGVQLADHY QQNTPIGDGP VLLPDNHYLS YQSKLSKDPN EKRDHMVLLE

             101                   121                          150
Tq-Ca-FLITS  FVTAAGITLG MDELYQGGSG GMVSKGEELF TGVVPILVEL DGDVNGHKFS
G-Ca-FLITS   FVTAAGITLG MDELYQGGSG GMVSKGEELF TGVVPILVEL DGDVNGHKFS

             151                   171                          200
Tq-Ca-FLITS  VSGEGEGDAT YGKLTLKFIC TTGKLPVPWP TLVTTLSWGV QCFARYPDHM
G-Ca-FLITS   VSGEGEGDAT YGKLTLKFIC TTGKLPVPWP TLVTTLSWGV QCFARYPDHM

             201                   221                          250
Tq-Ca-FLITS  KQHDFFKSAM PEGYVQERTI FFKDDGNYKT RAEVKFEGDT LVNRIELKGI
G-Ca-FLITS   KQHDFFKSAM PEGYVQERTI FFKDDGNYKT RAEVKFEGDT LVNRIELKGI

             251                   271                          300
Tq-Ca-FLITS  DFKEDGNILG HKLEYNYYSD NTRDQLTEEQ IAEFKEAFSL FDKDGDGTIT
G-Ca-FLITS   DFKEDGNILG HKLEYNYYSD DTRDQLTEEQ IAEFKEAFSL FDKDGDGTIT

             301                   321                          350
Tq-Ca-FLITS  TKELGTVMRS LGQNPTEAEL QDMINEVDAD GDGTFDFPEF LTMMARKMND
G-Ca-FLITS   TKELGTVMRS LGQNPTEAEL QDMINEVDAD GDGTFDFPEF LTMMARKMND

             351                   371                          400
Tq-Ca-FLITS  TDSEEEIREA FRVFDKDGNG YIGAAELRHV MTDLGEKLTD EEVDEMIRVA
G-Ca-FLITS   TDSEEEIREA FRVFDKDGNG YIGAAELRHV MTDLGEKLTD EEVDEMIRVA

             401               420
Tq-Ca-FLITS  DIDGDGQVNY EEFVQMMTAK
G-Ca-FLITS   DIDGDGQVNY EEFVQMMTAK
```

**Figure S4**. **Sequence alignment of Tq-Ca-FLITS and G-Ca-FLITS**. Residues with **yellow** background highlight mutations in G-Ca-FLITS relative to the parental Tq-Ca-FLITS. The RS20 peptide is in **blue**, the FP in **black**, and calmodulin in **orange**. Linker residues are **underlined**.

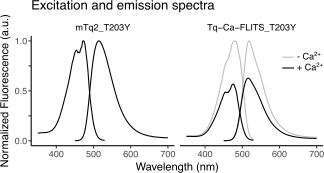

**Figure S5. Excitation and emission spectra of Tq-Ca-FLITS_T203Y and mTq2_T203Y.** The gray line indicates the calcium-free state (10 mM EGTA) and the black line the calcium-bound state (39 μM free Ca^2+^). Spectra are normalized to the top of the calcium-free state.

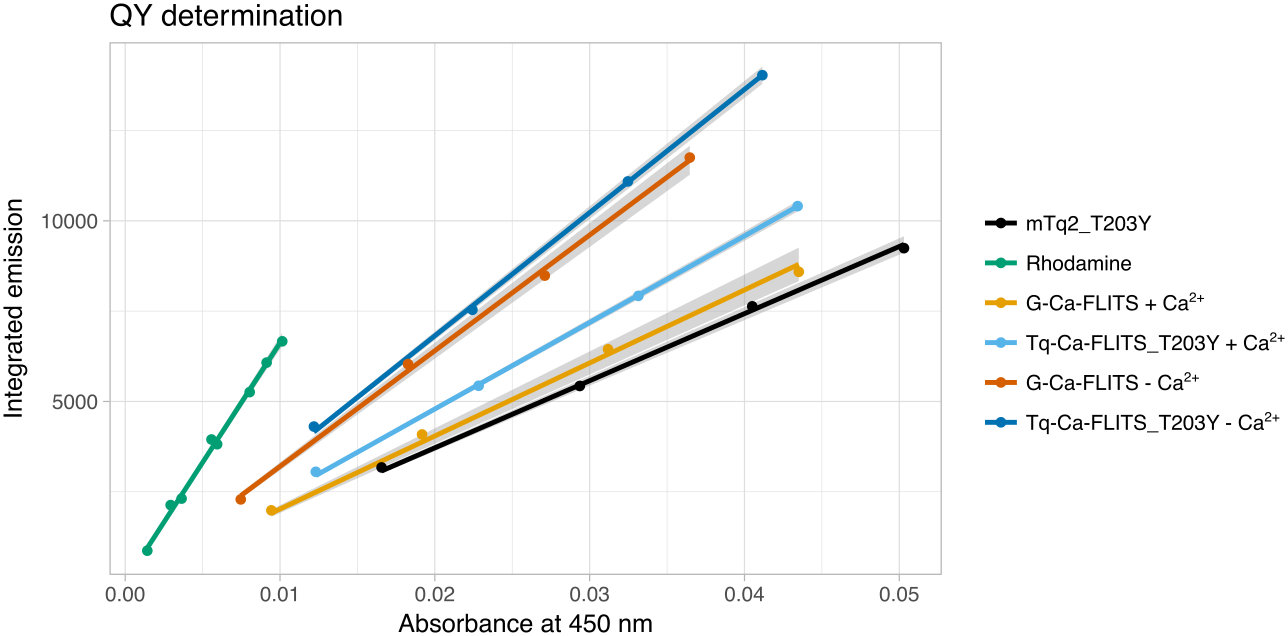

**Figure S6. QY determination.** For each protein, the absorbance and emission spectra were measured of four dilutions (indicated by points). This was done in both presence (39 μM free Ca^2+^) and absence of calcium (10 mM EGTA) for the sensor variants. For rhodamine, we used eight dilutions on two days. The measured absorbance and the integrated emission are plotted here. Linear models (lines) are fitted through the measurements and forced through the origin. Gray bands indicate the 95% confidence interval of the models.

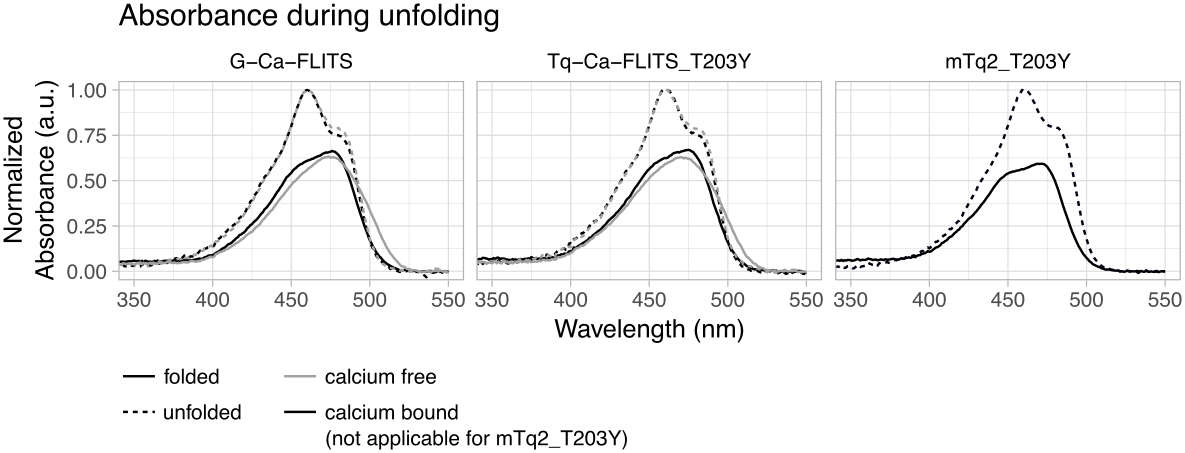

**Figure S7. Determination of the extinction coefficient by unfolding of the proteins.** Absorbance was measured before (solid line) and after (dotted line) unfolding by 0.5 M NaOH and 2 M urea of the proteins G-Ca-FLITS, Tq-Ca-FLTIS_T203Y and mTq2_T203Y. For both sensors, the determination was done in presence (black, 39 μM free Ca^2+^) and absence (gray, 10 mM EGTA) of calcium. Spectra are corrected for difference in dilution between the folded and unfolded state and normalized tot the maximum of the unfolded spectrum.

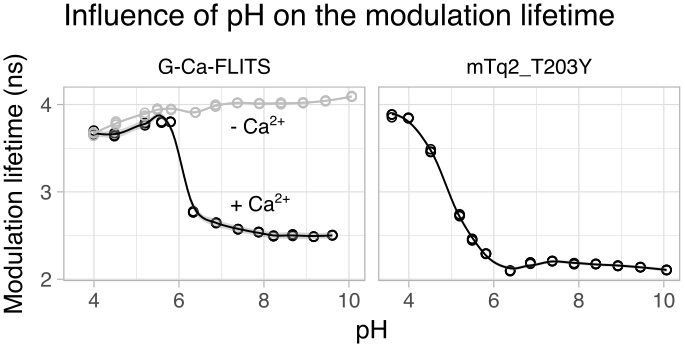

**Figure S8. Influence of pH on modulation lifetime of the proteins.** Fluorescence lifetime of proteins diluted in pH buffer was measured (n=3). In case of G-Ca-FLITS, this was done in presence (gray, 0.1 mM CaCl_2_) or absence of calcium (black, 5 mM EGTA). A smooth curve is fitted through the data using the loess method, using α = 0.4. The gray band indicated the 95% confidence interval of the smooth fit.

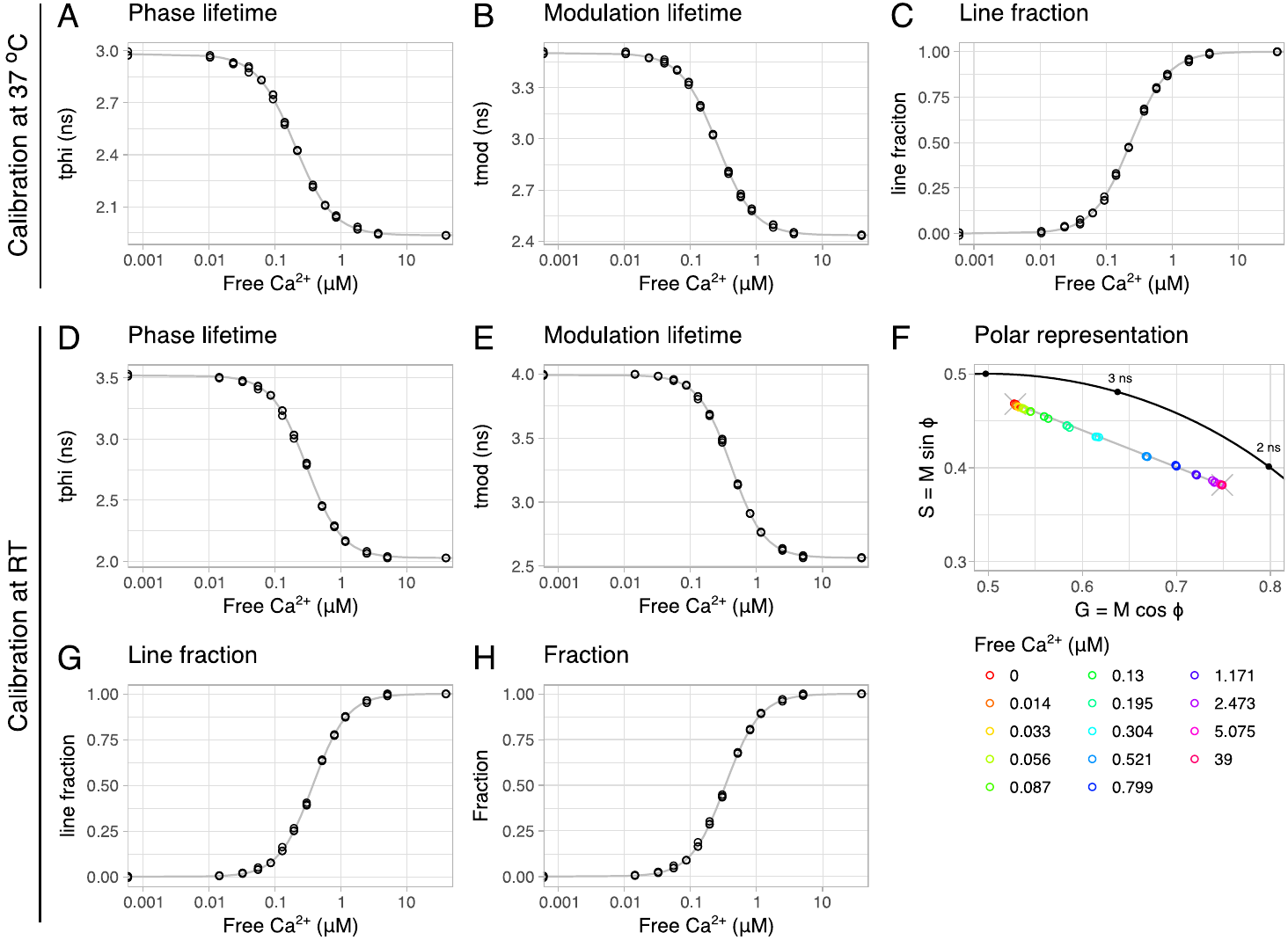

**Figure S9. Calcium calibration of G-Ca-FLITS in vitro at 37°C and at RT**. (**A**)Measured fluorescent phase and (**B**) modulation lifetime at 37°C for a range of calcium concentrations. (**C**) The line fraction of sensors in the calcium-bound state (the low lifetime) is determined, without considering the intensity contributions of the states, Kd = 234 nM and Hill coefficient = 1.5. (**D**) Measured fluorescent phase and (**E**) modulation lifetime at RT for a range of calcium concentrations. (**F**) The fluorescence lifetime at RT is plotted in a polar space, showing a straight line (in gray) between the average of the lowest and highest concentration (indicated by an X). (**G**) The line fraction is determined for each measurement at RT, with respect to the calcium-bound state (low lifetime state), Kd = 337 nM and Hill coefficient = 1.7. (**H**) The true fraction of sensors in the calcium-bound state at RT is determined by considering the intensity contribution of both states. In each panel the circles depict the individual measurements (n=3) and the gray lines are the fitted calibration curves. In the polar plot the concentration of free calcium is indicated by the color of the circles.

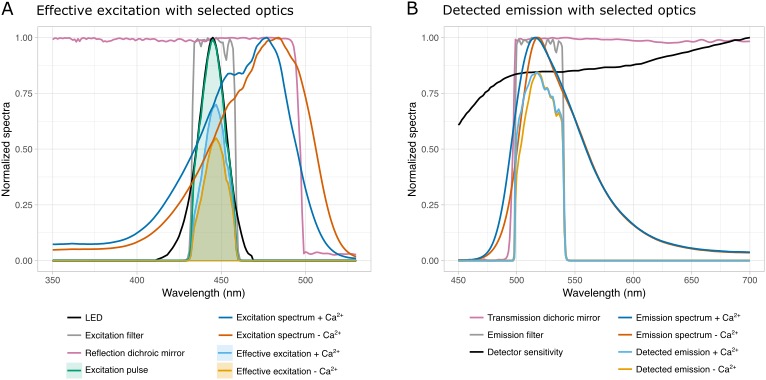

**Figure S10. Effective excitation and detected emission of G-Ca-FLITS for the indicated filters.** Excitation and emission spectra were recorded using isolated protein. Filter profiles and the detector sensitivity are provided on the manufacturers’ respective websites. The LED brightness was measured**.** (**A**) All normalized spectra related to excitation of G-Ca-FLITS are shown. The “effective excitation pulse” (“LED” $\times$ **“**excitation filter” $\times$ “reflection dichroic mirror”) is shown in green. The “effective excitation pulse” is multiplied with the “excitation spectra” to yield the “effective excitation” of G-Ca-FLITS for each state (cyan and orange). The $e_{rel}$ discussed in Supplemental Note 1, is the integral of “effective excitation” divided over de integral of “excitation pulse”. (**B**) All normalized spectra related to emission and detection of G-Ca-FLITS are shown. The “detected emission” of G-Ca-FLITS was calculated for each state by multiplication of the following spectra: “transmission dichroic mirror” $\times$ **“**emission filter” $\times$ “detector sensitivity” $\times$ “emission spectrum”. The $S_{rel}$ discussed in Supplemental Note 1, is the integral of “detected emission” divided over de integral of “emission spectra” for each calcium state.

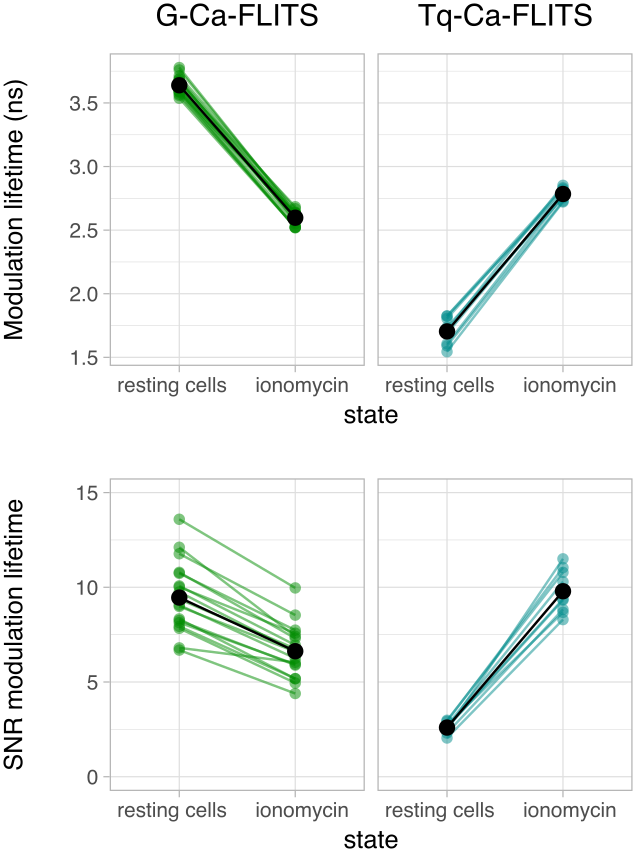

**Figure S11. Comparison of the signal-to-noise ratio of the modulation lifetime of G-Ca-FLITS and Tq-Ca-FLITS in HeLa cells.** HeLa cells were measured in a resting state and after addition of 5 μg/ml ionomycin and 5 mM calcium. Measurements of individual cells are indicated by colored dots (n=18 for G-Ca-FLITS, n=9 for Tq-Ca-FLITS), averages of all cells are indicated in black. For each cell, 400 pixels were analyzed for the mean lifetime and the sd of the lifetime. SNR = mean lifetime / sd lifetime. Comparable modulation lifetimes are found for all cells expressing a construct, however, the SNR is lower for lower intensity cells and varies less between the two states for G-Ca-FLITS.

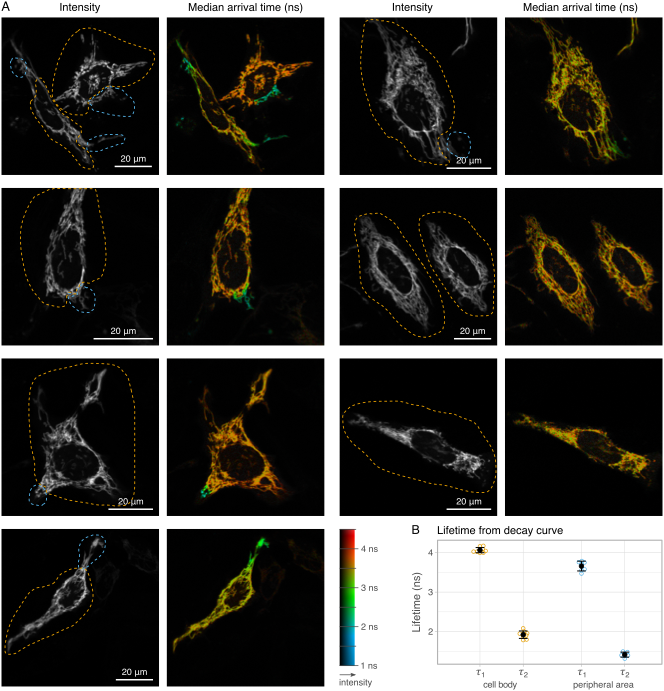

**Figure S12. Confocal lifetime imaging of G-Ca-FLITS expressed in mitochondria of HeLa cells.** (**A**) Intensity and false color images of the median arrival time. The peripheral regions with a differential lifetime are indicated in blue, the cell bodies are indicated in orange. Images were taken with a PicoQuant TCSPC setup with 60x magnification with a total integration time of ~ 5 min. (**B**) Two lifetime components (𝜏_1_ and 𝜏_2_) determined from decay curves of the two types of areas. Each circle indicates either a single ‘peripheral area’ (blue) or a single ‘cell body’ (orange). The mean is indicated in black, including the standard deviation

**A**

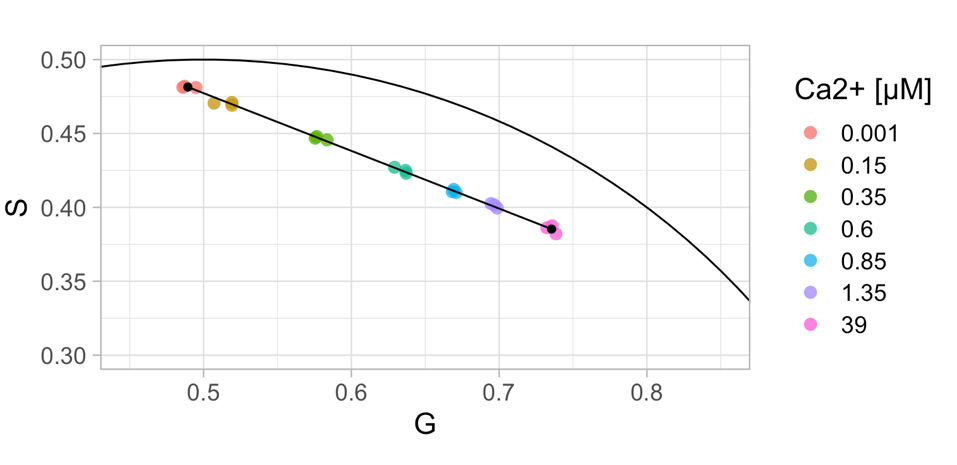

**B**

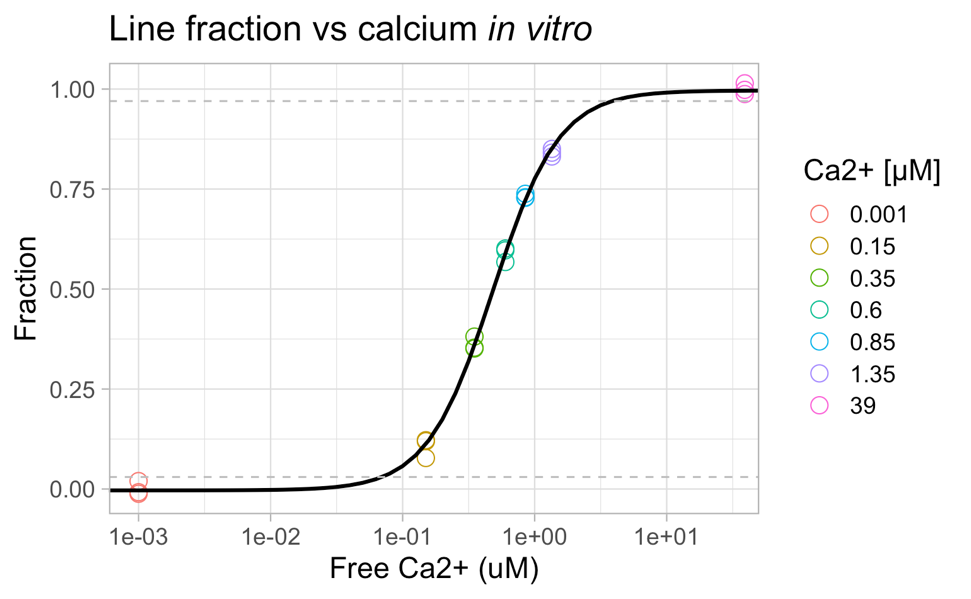

**Figure S13. In vitro calibration of G-Ca-FLITS on the Leica Stellaris8 at room temperature.** (**A**) Purified protein was added to calcium buffers and lifetimes images were acquired at room temperature. The fluorescence lifetime in a range of calcium concentrations is plotted in a polar space (n=3), with the color indicating the concentration. The measurements fall on a straight line on the polar plot between the lowest and highest concentration. (**B**) For each measurement, the fraction of sensors in the calcium-bound state (the low lifetime) is determined, taking the intensity contribution of the two extreme states into account. The fraction is plotted against the concentration of free calcium to obtain a calibration curve. The solid line is a fit with Kd = 0.483 µM and Hill coefficient 1.7.

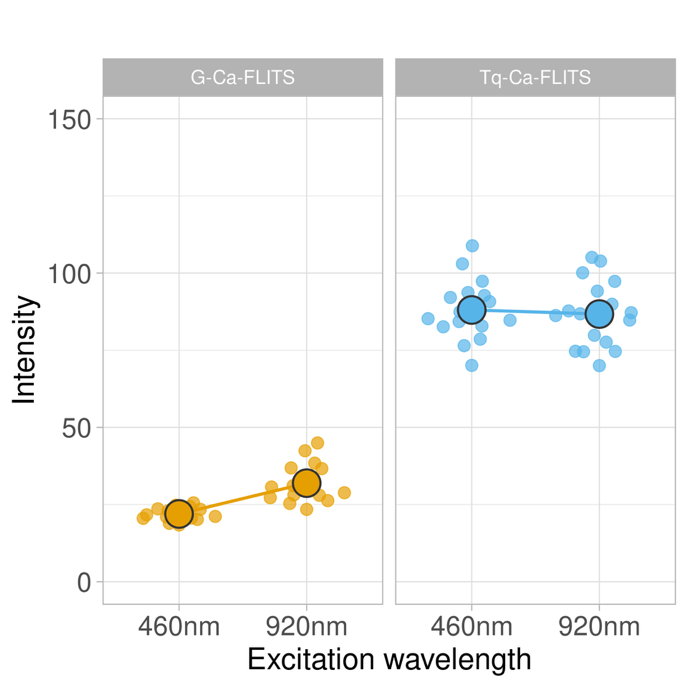

**Figure S14. Similar brightness for G-Ca-FLITS and Tq-Ca-FLITS between 1PE and 2PE.** Ni-NTA beads were loaded with purified Tq-Ca-FLITS or G-Ca-FLITS that are both equipped with a 6xHis tag. Images were acquired with either 1PE (460 nm) or 2PE (920 nm) and the intensities of the beads were quantified. The smaller dots are average intensity of individual beads and the larger dot is the average. The overall higher intensity of Tq-Ca-FLITS is due to a higher concentration of protein loaded on the beads.

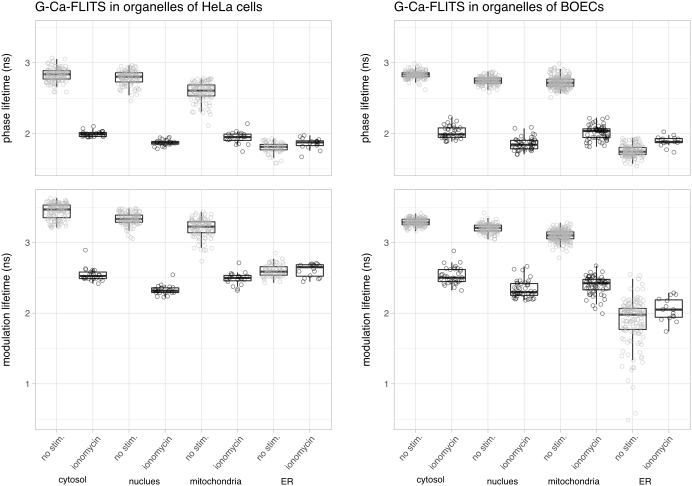

**Figure S15. Aggregated FLIM data of measurements in organelles under resting conditions and after incubation with ionomycin.**

**Table S1. Lifetime and spectral screen of candidate sensors to create G-Ca-FLITS.** Sensor variants are indicated by their circular permutation and mutations in the FP in the sensor. For example, ‘cp146’ has from N- to C-terminus the following domains: calmodulin binding peptide M13, amino acids 146-283 of mTq2, a flexible GGSGG linker, amino acids 1-145 of mTq2, CaM (calmodulin). ‘T203Y’ indicates a T203Y mutation on the mTq2. The calcium-bound state is measured in presence of 0.1 mM CaCl_2_ and the calcium-free state after addition of 9.5 mM EDTA.

*Emission and excitation maxima are only indicated if a red shift of the spectrum with respect to Tq-Ca-FLITS was observed. See also Figure S3.

**Phase lifetimes were measured at 37 °C by FD-FLIM. The lifetime change is calculated as the phase lifetime in the calcium-bound state minus the calcium-free state.

| **Sensor variant** | **Emission maximum (nm)*** | | **Excitation maximum (nm)*** | | **Phase lifetime (ns) **** | | |
| --- | --- | --- | --- | --- | --- | --- | --- |
|  | **calcium free** | **calcium bound** | **calcium free** | **calcium bound** | **calcium free** | **calcium bound** | **change** |
| **Tq-Ca-FLITS** | 492, 502 | 480, 507 | 437, 455 | 437, 456 | 1.72 | 2.86 | 1.14 |
| **T203Y** | 516 | 514 | 461, 488 | 459, 478 | 3.08 | 2.20 | -0.88 |
| **cp146, T203Y** | 518 | 517 | 461, 480 | 460, 479 | 3.40 | 3.13 | -0.28 |
| **cp147, T203Y** | 517 | 518 | 459, 480 | 460, 481 | 3.02 | 3.38 | 0.36 |
| **cp149, T203Y** | 516 | 518 | 461, 481 | 460, 479 | 2.69 | 2.65 | -0.04 |
| **cp150, T203Y** | 512, shoulder 490 | 515 | 460, shoulder 435 | 460, 478 | 1.85 | 2.37 | 0.52 |
| **cp151, T203Y** | 516 | 516 | 460, 479 | 460, 479 | 2.73 | 2.91 | 0.18 |
| **cp152, T203Y** | 516 | 517 | 460, 480 | 460, 479 | 3.16 | 3.05 | -0.11 |
| **cp146, T203H** | no red shift | no red shift | no red shift | no red shift | 3.09 | 3.21 | 0.12 |
| **cp147, T203H** | no red shift | no red shift | no red shift | no red shift | 3.13 | 2.86 | -0.27 |
| **cp149, T203H** | no red shift | no red shift | no red shift | no red shift | 2.67 | 2.78 | 0.11 |
| **cp150, T203H** | no red shift | no red shift | no red shift | no red shift | 2.60 | 2.96 | 0.36 |
| **cp152, T203H** | no red shift | no red shift | no red shift | no red shift | 3.15 | 3.01 | -0.13 |

**Table S2.** All primers were ordered from Integrated DNA Technologies. Mixed bases are indicated by special letters: Y (C or T), R (A or G), K (G or T), M (A or C), D (A, G or T), H (A, C or T) and N (any).

| No. | Sequence | Internal name |
| --- | --- | --- |
| *Engineering G-Ca-FLITS* | |  |
| 1 | CAACCACTACCTGAGCYACCAGTCCAAGCTGAGCAAAGAC | FW_Tq2_T203Y/H |
| 2 | GTRGCTCAGGTAGTGGTTGTCGG | RV_Tq2_T203Y/H |
| 3 | CAACCACTACCTGAGCTTCCAGTCCAAGCTGAGCAAAGAC | FW_Tq2_T203F |
| 4 | GAAGCTCAGGTAGTGGTTGTCGG | RV_Tq2_T203F |
| 5 | CAAGGCCAACTTCAAGYDSCGCCACAACATCGAGGAC | Fw_Tq2_I167arom |
| 6 | SHRCTTGAAGTTGGCCTTGATGCCG | Rv_Tq2_I167arom |
| 7 | GCTGAGCTCACCCGTGNNKGTCTATATCACCGCCG | FW_G-FLITS_PVX-linker |
| 8 | GTCACGCGTMNNGTCGCTATAGTAGTTGTAC | RV_G-FLITS_SDX-end |
| *Creation dual expression ratio plasmid* | |  |
| 9 | CGGCATGGACGAGCTGTACAAGTCGACCAG | FW-del_SacI_in-FP-pDress |
| 10 | CTGGTCGACTTGTACAGCTCGTCCATGCCG | RV-del_SacI_in-FP-pDress |
| 11 | GTCGCGAATTCAGGCGC | FW_FP-lessFRET-P2A |
| 12 | TTTGGATCCAGCGCTAGCGAAGGTCC | RV_FP-lessFRET-P2A |
| 13 | TAAGGATCCCGCCACAATGGTC | FW_FLITS-backbone_notags |
| 14 | GCGCCTGAATTCGCGAC | RV_FLITS_backbone_notags |
| *Sensors and FPs in ratioplasmid* | |  |
| 15 | AACCACTACCTGAGCTACCAGTCCAAGCTGAGC | FW_Tq2_T203Y |
| 16 | GCTCAGCTTGGACTGGTAGCTCAGGTAGTGGTT | RV_Tq2_T203Y |
| 17 | GCTGGATCCCGCCACCATGGTGAGC | FW_FP_to_pFR |
| 18 | CACAAGCTTTTACTTGTACAGCTCGTCC | RV_FP_to_pFR |
| 19 | GCTGGATCCCGCCACA ATGGTCGACTCATCACG | FW_RCaMPs_to_ratio |
| 20 | CACAAGCTTCTACTTCGCTGTCATCATTTG | RV_RCaMPs_to_ratio |
| *Organelle targeting* | |  |
| 21 | GCTACCGGTCGCCACCATGGTCGACTCTTCACG | FW_VY_to_C1N1 |
| 22 | TTTTGTACACCTTCGCTGTCATCATTTGGACAAACTC | RV_VY_to_C1N1 |
| 23 | TTTTGTACACCTACTTCGCTGTCATCATTTGGACAAACTC | RV_VY-STOP_to_C1N1 |
